# From static thresholds to dynamic waves: How biological memory destabilizes malaria transmission Potential

**DOI:** 10.64898/2026.05.11.724460

**Authors:** Steeven B. Affognon, Priscille Barreaux, Shirley Abelman, Antoine M. G. Barreaux

## Abstract

The basic reproduction number *R*_0_ is central to malaria epidemiology, yet it is typically treated as a static quantity derived under memoryless assumptions for mosquito demography. In natural systems, however, mosquito populations are shaped by delayed processes such as larval development and density-dependent feedback, introducing biological memory into vector dynamics. We develop a minimal delay-based framework that incorporates this memory into the Ross–Macdonald model by describing adult mosquito abundance with a retarded differential equation. This formulation induces a time-dependent transmission potential *R*_0_(*t*). Using complex analysis and the argument principle, we derive an explicit stability threshold 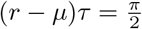, which separates stable from oscillatory transmission regimes. Near this threshold, delayed feedback produces slow relaxation times and sustained transient oscillations, implying that transmission potential may vary intrinsically even in the absence of external forcing. To account for ecological variability, we extend this deterministic condition into a probabilistic framework and define the stability probability as 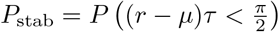. Numerical simulations and global sensitivity analysis show that recruitment and developmental delays are the primary drivers of instability, while adult mortality has a weaker stabilizing effect. These results indicate that malaria interventions may influence not only the magnitude of malaria transmission but also its dynamical stability. By linking delay dynamics, transmission theory, and uncertainty quantification, this framework provides a basis for stability-aware modeling and interpretation of malaria transmission under ecological variability.

**Author summary:** Malaria transmission is often summarized by a single number, *R*_0_, treated as a fixed indicator of whether transmission will increase or decline. This assumes mosquito populations respond instantly to environmental conditions. In reality, mosquitoes develop through stages where larval conditions, such as crowding, nutrition, or temperature, affect adult populations only after a delay. This creates biological memory: today’s mosquitoes reflect past environments. We show that this memory can fundamentally reshape transmission dynamics. When developmental delays are included, transmission potential is no longer constant but can fluctuate over time, even in stable environments. These fluctuations can persist or amplify depending on the balance between mosquito growth, mortality, and delay. As a result, variability in mosquito abundance or malaria transmission may arise from intrinsic dynamics rather than external drivers alone. Under ecological variability, stability becomes probabilistic, allowing estimation of how likely transmission is to remain stable. Interventions that reduce larval productivity or increase adult mortality may therefore both lower transmission and make it more predictable, improving interpretation and control strategies.

## Introduction

The basic reproduction number *R*_0_ is a cornerstone of infectious disease modeling and, in the context of vector-borne pathogens, represents the standard measure of transmission potential in a host–vector system [1–4]. In the classical Ross–Macdonald framework, *R*_0_ is derived from a suite of entomological and epidemiological parameters, including mosquito density, biting rate, mosquito survival, and parasite development, and has long provided the foundation for malaria theory and control design [1–5].However, *R*_0_ is typically treated as a static threshold quantity, implicitly assuming memoryless mosquito demography in which population regulation responds instantaneously to environmental conditions. Under this assumption, mosquito abundance is governed by fixed parameters, and transmission potential is interpreted as an invariant property of the system [6–8].

In natural ecosystems, this assumption is rarely satisfied. Mosquito populations are structured by multistage life cycles spanning aquatic and terrestrial environments, where larval development, stage progression, and density-dependent competition introduce intrinsic temporal delays [9–11]. These processes generate biological memory, such as adult abundance reflects past ecological conditions rather than only current states. Within this developmental framework, mosquito fitness is shaped by sensitive periods during larval development, when environmental conditions such as temperature, nutrition, and crowding determine adult traits [12–14]. These carry-over effects influence into traits directly relevant for transmission, including body size, longevity, and vector competence [15, 16]. In addition, maternal environmental effects can propagate across generations, where exposure to pathogens or insecticides alters offspring development or immune priming [13, 17–19].

Together, these mechanisms generate delayed density dependence in mosquito populations, a well-known driver of oscillatory and non-equilibrium dynamics in ecological systems [9–11, 20]. As a result, mosquito abundance cannot generally be understood as an instantaneous response to current conditions alone, but instead reflects integrated past ecological forcing [6–8]. In malaria systems, such intrinsic delays provide a mechanistic explanation for temporal variability in vector abundance and transmission intensity that is not fully captured by seasonal forcing alone [21–23].Because *R*_0_ depends directly on mosquito abundance and life-history traits, delayed feedback can affect not only equilibrium transmission intensity but also its temporal stability and predictability [24, 25]. Importantly, these delays also interact with the parasite development: the extrinsic incubation period (EIP) depends on mosquito age, nutrition, and developmental history. Given that EIP and mosquito longevity are often of comparable magnitude, small shifts in developmental timing can produce strongly nonlinear effects in the fraction of mosquitoes surviving to infectiousness [12, 17].

Mathematically, delay differential equations provide a natural framework for representing biological memory [11, 20, 26]. Unlike age-structured or stage-structured partial differential equation models, which require full demographic resolution, delay differential equations allow developmental delay to be isolated as a mechanistic parameter governing feedback between past and present population states. This makes them particularly suitable for analyzing how temporal lags influence transmission stability while maintaining analytical tractability. Previous work has incorporated delays into malaria and vector-borne disease models from multiple perspectives, including parasite incubation dynamics [27, 28], vaccination and waning immunity [29, 30], and infection progression or treatment dynamics [31, 32]. However, these studies primarily focus on disease prevalence, equilibrium stability, or coupled host–vector dynamics, leaving the probabilistic interpretation of transmission thresholds under demographic memory less explicitly characterized.

By contrast, the present study addresses a more fundamental question: how does delayed mosquito demography reshape the temporal behavior, stability, and interpretation of the transmission potential itself? This distinction is important because treating *R*_0_ as static may conceal dynamical instabilities driven by biological memory, limiting the interpretation of vector control strategies. Moreover, mosquito populations are often most sensitive during specific developmental windows, meaning that transient environmental perturbations can propagate through delayed feedback loops and produce long-lasting effects on system dynamics [12, 13, 15]. These features naturally motivate an uncertainty-aware perspective, in which demographic variability is explicitly incorporated into transmission stability. Here, we develop a minimal delay-based framework that embeds mosquito demographic memory directly into malaria transmission theory. Adult mosquito abundance is modeled using a retarded differential equation with delayed density dependence, producing a time-dependent transmission potential within the classical Ross–Macdonald structure. This shifts *R*_0_ from a static threshold to a dynamical object shaped by ecological feedback over time.

We address three main questions: (i) how delayed mosquito demography affects the local stability of transmission equilibria; (ii) under what conditions biological memory can generate intrinsic temporal variability in transmission potential in the absence of external forcing; and (iii) how stability can be interpreted probabilistically under ecological uncertainty in demographic parameters. We derive an explicit stability threshold using complex analysis, characterize delay-induced oscillatory regimes, and extend the deterministic condition into a probabilistic framework for stability under variability. Overall, this work shows that biological memory in mosquito demography fundamentally reshapes not only the magnitude of malaria transmission potential, but also its stability structure and predictability, providing a theoretical basis for stability-aware interpretation of vector control strategies.

## Materials and methods

As a conceptual overview Fig 1, we present a progression from classical static transmission thresholds to delay-driven dynamics and finally to probabilistic stability regimes. This framework reflects increasing biological realism in how mosquito populations are represented, moving from instantaneous equilibrium assumptions to models that incorporate intrinsic temporal memory and uncertainty in demographic parameters. A key implication is that malaria transmission is shaped not only by the magnitude of vector abundance, but also by the timing of demographic responses relative to the memory timescale. Consequently, interventions that modify mosquito life-history traits, such as survival, developmental time or reproduction, may influence both the mean level of transmission and its temporal stability with implications for predictability and control performance.

**Fig 1.**
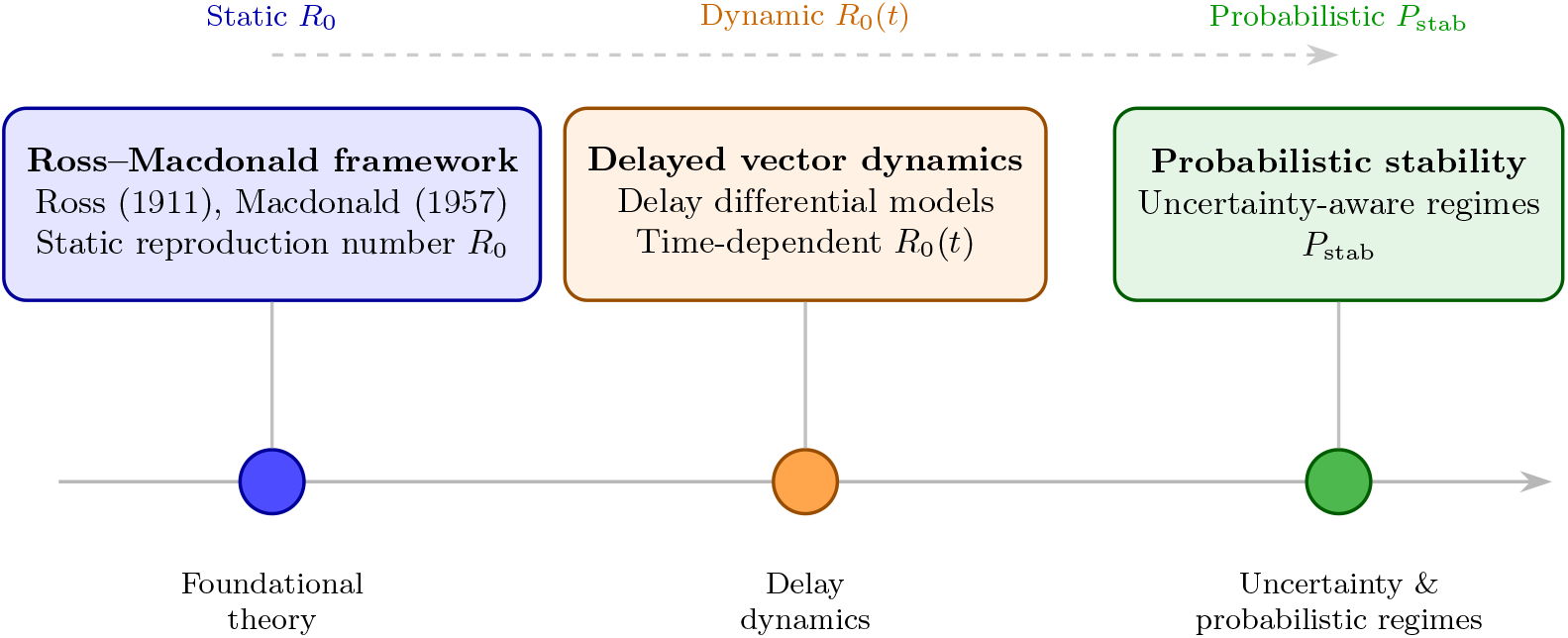
Conceptual progression of malaria transmission modeling. Classical Ross–Macdonald theory defines the basic reproduction number *R*_0_ as a static measure of transmission potential [1, 2]. Subsequent extensions incorporated delayed demographic processes using delay differential equations, producing time-dependent transmission dynamics. In this study, we further extend this framework by interpreting delay-induced stability thresholds in probabilistic terms, yielding a stability probability *P*_stab_ under ecological and demographic uncertainty. This progression highlights a shift from static thresholds to deterministic delay dynamics, and finally to uncertainty-aware stability regimes.

### Delayed mosquito demography

Let *x*(*t*) denote the abundance of adult female mosquitoes at time *t*. We model mosquito population dynamics using a delay differential equation that incorporates maturation time and delayed density dependence:

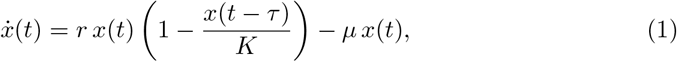

where *r >* 0 is the intrinsic recruitment rate, *K >* 0 is the carrying capacity, *µ >* 0 is the adult mosquito mortality rate, and *τ >* 0 is a discrete delay representing the developmental time from larval recruitment to adult emergence, as well as aggregate density-dependent maturation effects. This formulation assumes that current adult growth is regulated by past population density, reflecting the finite time required for immature stages to contribute to the adult population. As such, *τ* encodes biological memory arising from stage-structured development without explicitly resolving juvenile dynamics.

#### History function and biological interpretation

Equation (1) requires specification of a history function *x*(*t*) = *φ*(*t*) for *t* ∈ [−*τ*, 0]. Mathematically, the state of the system belongs to the function space *C*([−*τ*, 0]), reflecting that the future evolution depends on the recent past rather than a single initial state [11]. Biologically, *φ* represents the pre-existing adult female population over the delay interval, which seeds the current dynamics through delayed density-dependent recruitment. In this framework, the history is not interpreted as a detailed mechanistic record of all life stages, but rather as an effective summary of prior adult abundance that has already undergone immature-stage processing. In other words, the system “inherits” memory through the adult population trajectory over the interval [−*τ*, 0], which implicitly encodes past oviposition, larval survival, and maturation outcomes. This interpretation is particularly relevant in ecological systems where developmental processes introduce unavoidable lag between cause and effect. The history function therefore captures the biological reality that current adult abundance is shaped by conditions experienced over a finite developmental window, rather than by instantaneous feedback alone.

#### Modeling assumptions and biological scope

Equation (1) is formulated as a reduced adult-population model in which delayed feedback represents the aggregate effect of past immature-stage processes on current adult growth. In this formulation, the recruitment parameter *r* is an effective demographic rate that combines oviposition, immature survival, and successful emergence into the adult female population. Consequently, larval and pupal stages are not represented as explicit state variables. The delay *τ* is treated as a constant effective ecological memory term. Biologically, it represents the time required for cohorts influenced by past density conditions to contribute to the adult population. This includes development duration, but also indirect effects such as competition-induced delays in maturation. While real mosquito development time is environmentally heterogeneous, varying with temperature, nutrition, and larval density, the present formulation assumes a time-invariant approximation to isolate the role of memory in shaping population stability. This simplification is intentional. By collapsing stage-structured dynamics into a single delayed feedback term, the model isolates the qualitative consequences of biological memory on mosquito population regulation. In particular, it allows one to disentangle how delay alone can generate oscillatory behavior, alter stability boundaries, and reshape long-term population trajectories. Extensions of this framework are straightforward. A more mechanistic representation could introduce a state-dependent delay *τ* = *τ* (*x*), allowing developmental time to vary with density or environmental conditions. Alternatively, a multi-compartment model with explicit larval dynamics could be used to resolve immature-stage processes directly. However, such refinements are deferred in this study in favor of analytical tractability and clarity in identifying delay-driven stability effects. Overall, the present model should be viewed as a minimal mechanistic representation that preserves the essential feature of interest, temporal memory in demographic feedback, while remaining sufficiently simple to permit analytical insight into stability and transmission-relevant dynamics

#### Method of steps

Solutions of (1) can be constructed interval by interval using the method of steps [20]. On the first interval [0, *τ*], the delayed term *x*(*t*−*τ*) is entirely determined by the history function *φ*. On the subsequent interval [*τ*, 2*τ*], the delayed term is governed by the solution previously computed on [0, *τ*], and the procedure then continues recursively. This construction possesses a natural cohort-based interpretation in mosquito demography: each step in the solution corresponds to a successive generation of adults whose recruitment is regulated by the density of the specific cohort that preceded them by one developmental window.

### Transmission potential as a dynamical quantity

The classical Ross–Macdonald expression for malaria transmission potential is traditionally defined as

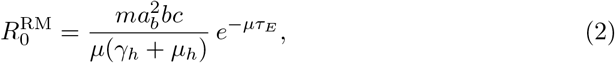

where *m* = *N*_*v*_*/N*_*h*_ is the mosquito-to-human ratio, *a*_*b*_ is the biting rate, *b* and *c* are transmission probabilities, *γ*_*h*_ is the human recovery rate, *µ*_*h*_ is the human mortality rate, and *τ*_*E*_ is the extrinsic incubation period [3]. In this reduced framework, we allow mosquito abundance to vary dynamically while treating the epidemiological parameters as an aggregate constant *C*_0_. This assumes that biting rate, transmission probabilities, and the EIP do not vary with mosquito density or temperature, standard simplifying assumptions used here to isolate the demographic feedback.

Given *m*(*t*) = *x*(*t*)*/N*_*h*_, we define the dynamic transmission potential:

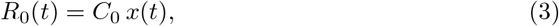

With

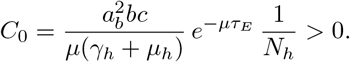

Under this definition, *R*_0_(*t*) is directly proportional to adult mosquito abundance. Substituting this relationship into the demographic model (1) yields

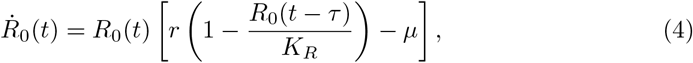

where *K*_*R*_ = *C*_0_*K*. Hence, the biological memory inherent in mosquito demography induces a corresponding memory in the transmission potential of the system.

### Equilibria and linearization

We rewrite (1) in the standard functional form

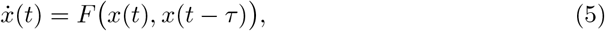

where the bivariate function *F* : ℝ^2^→ ℝ is given by

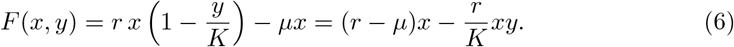

Here, *x* denotes the current adult mosquito density and *y* the delayed density. Introducing *F* (*x, y*) makes it possible to linearize the system using partial derivatives with respect to the present and delayed arguments, as is standard for retarded functional differential equations [11, 20].

#### Equilibria

An equilibrium of (1) is a constant solution *x*(*t*) ≡*x*\*, for which *x*(*t*−*τ*) ≡*x*\* as well. Substituting into (5) gives

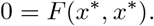

Using (6), this becomes

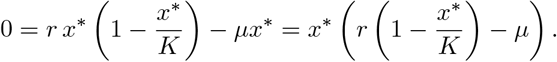

Hence there are two equilibria:

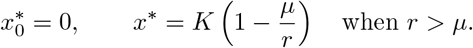

The equilibrium *x*\* is biologically meaningful only when *r > µ*, and it represents the positive adult mosquito population maintained by the balance between recruitment, mortality, and delayed density dependence.

Since *R*_0_(*t*) = *C*_0_*x*(*t*) with *C*_0_ *>* 0, the corresponding equilibrium transmission potential is

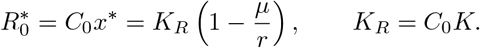

#### Linearization about the positive equilibrium

To study local stability of the positive equilibrium, we introduce the perturbation

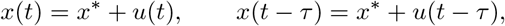

where *u*(*t*) is assumed to be small. Substituting into (5) and retaining only first-order terms yields

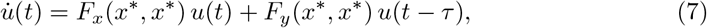

where *F*_*x*_ and *F*_*y*_ denote partial derivatives of *F* with respect to its first and second arguments.

From (6),

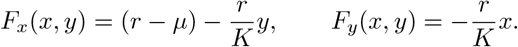

Evaluating at (*x*\*, *x*\*), and using *x*\* = *K*(1 − *µ/r*), gives

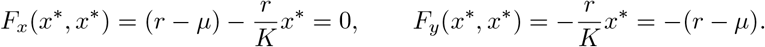

Therefore, the linearized delay differential equation becomes

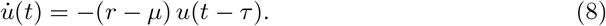

#### Characteristic equation

To determine local stability, we seek exponential solutions of the form *u*(*t*) = *e*^*λt*^, where *λ* ∈ ℂ. Substituting this ansatz into (8) gives

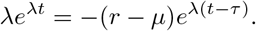

Dividing by *e*^*λt*^≠ 0 yields the characteristic equation

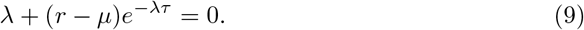

Because of the delay term *e*^−*λτ*^, (9) is transcendental and has infinitely many roots, generally complex. Local asymptotic stability of *x*\* holds if and only if all characteristic roots satisfy ℜ(*λ*) *<* 0 [11, 20, 26].

### Local stability analysis via complex analysis

The linearized equation (8) is a scalar retarded delay differential equation. Unlike ordinary differential equations, its stability cannot be determined from a finite-dimensional Jacobian matrix. Instead, stability is governed by the location of the roots of the characteristic equation

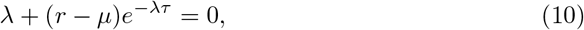

which is transcendental due to the exponential delay term.

#### Characteristic roots and their meaning

A characteristic root *λ* ∈ ℂ corresponds to a mode

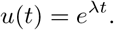

If ℜ(*λ*) *<* 0, the mode decays exponentially and contributes to stability. If ℜ(*λ*) *>* 0, the mode grows exponentially and destabilizes the equilibrium. Complex roots *λ* = *α*±*iω* generate oscillatory modes, reflecting the presence of delayed feedback in the system.

#### The characteristic function

Define the characteristic function

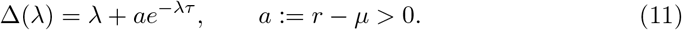

The function Δ is entire and has no poles. Its zeros coincide exactly with the characteristic roots of the linearized system.

#### Nyquist contour and root counting

Let *C*_*R*_ denote the contour consisting of:

- the line segment *λ* = *iω, ω* ∈ [−*R, R*],
- the semicircle *λ* = *Re*^*iθ*^, *θ* ∈ [−*π/*2, *π/*2], which encloses the right half-plane up to radius *R*.

Let *N*_*R*_ denote the number of zeros of Δ inside *C*_*R*_. Since Δ is entire, it has no poles. By the argument principle,

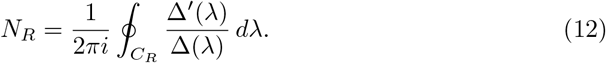

Equivalently, *N*_*R*_ equals the winding number of the curve Δ(*λ*), *λ* ∈ *C*_*R*_, around the origin.

#### Contribution of the large semicircle

On the semicircle *λ* = *Re*^*iθ*^, we have

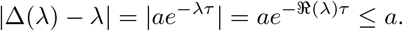

Since |*λ*| = *R* → ∞, it follows that

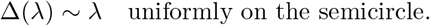

Thus, the contribution of the semicircle to the winding number vanishes as *R*→ ∞. The root count is therefore determined entirely by the image of the imaginary axis, known as the Nyquist curve.

#### Nyquist curve and root crossing

Setting *λ* = *iω*, we obtain

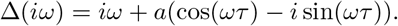

Hence

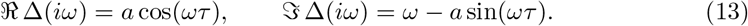

A change in stability occurs when a characteristic root crosses the imaginary axis, which corresponds to Δ(*iω*) = 0. Therefore,

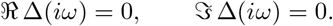

From (13), this yields

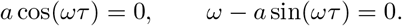

The first condition gives

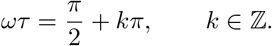

The smallest positive solution corresponds to *k* = 0, for which sin(*ωτ*) = 1. Substituting into the second equation gives

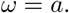

Hence the critical condition is

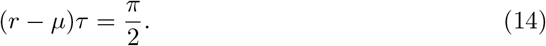

#### Stability conclusion

For (*r* − *µ*)*τ < π/*2, the Nyquist curve does not encircle the origin, so no characteristic roots lie in the right half-plane. For (*r*−*µ*)*τ > π/*2, the curve encircles the origin once, implying the existence of at least one root with positive real part.

##### Theorem 1

**(Delay-induced stability threshold)** *Let r > µ >* 0. *The positive equilibrium x*\* = *K*(1 − *µ/r*) *of* (1) *is locally asymptotically stable if*

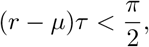

*and unstable if*

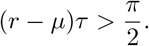

*Proof*. Stability is determined by the location of the roots of the characteristic equation (10). The argument principle shows that the number of roots in the right half-plane changes only when the Nyquist curve Δ(*iω*) crosses the origin. This occurs precisely when (*r*−*µ*)*τ* = *π/*2. For smaller values, no roots lie in the right half-plane, while for larger values, at least one root has positive real part. This proves the result. □

Near the critical boundary (*r*−*µ*)*τ* = *π/*2, the system is expected to exhibit slowly decaying oscillations on the stable side and emerging self-sustained oscillatory behavior on the unstable side, making this near-critical regime especially relevant for interpretation of transient dynamics and intervention response.

##### Proposition 1

**(Inheritance of stability by transmission potential)** *Let R*_0_(*t*) = *C*_0_*x*(*t*) *with C*_0_ *>* 0. *Then the equilibrium transmission potential* 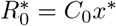 *is locally asymptotically stable if and only if the mosquito equilibrium x*\* *is locally asymptotically stable*.

*Proof*. The mapping *x* →*R*_0_ = *C*_0_*x* is linear, invertible, and positive. Therefore, perturbations of *x*\* and 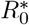 differ only by a constant scaling factor, and their stability properties coincide. □

### Probabilistic stability under ecological uncertainty

In realistic ecological environments, demographic parameters such as the recruitment rate *r*, mortality rate *µ*, and maturation delay *τ* are subject to environmental variability and spatial heterogeneity. We therefore model (*r, µ, τ*) as random variables defined on a probability space (Ω,ℱ, ℙ), with a joint distribution that may incorporate dependence between parameters.

#### Definition 1

**(Stability margin)** *Define the stability margin*

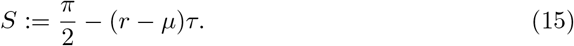

*The equilibrium is locally asymptotically stable when S >* 0 *and unstable when S <* 0.

#### Definition 2

**(Probability of stability)** *The probability of stability is defined by*

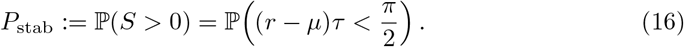

*The complementary probability*

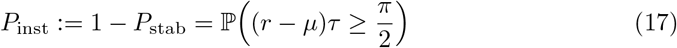

*can be interpreted as an instability risk index*.

#### Remark 1

**(Relation to classical** *R*_0_ *>* 1 **thresholding)** *Classical thresholding concerns invasion at the disease-free state. In contrast, P*_stab_ *characterizes the likelihood that the transmission potential approaches equilibrium without delay-induced oscillatory or unstable behavior. These perspectives are complementary but address distinct dynamical questions*.

Because *τ >* 0, (16) can equivalently be written as

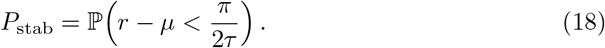

This representation highlights that stability becomes less likely as delays increase, and more likely as the net demographic response *r* − *µ* decreases.

#### Remark 2

**(Interpretation of the instability risk index)** *The quantity P*_inst_ = 1−*P*_stab_ *can be interpreted as an instability risk index, that is, the probability that delayed mosquito demography places the system in a regime where equilibrium transmission potential is not robustly stable. In applied terms, high values of P*_inst_ *indicate increased risk of endogenous oscillatory or poorly predictable transmission dynamics under ecological uncertainty*.

#### Proposition 2

**(Monte Carlo estimator of stability probability)** *Let* 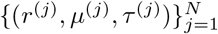 *be independent and identically distributed samples drawn from the joint distribution of* (*r, µ, τ*). *Define the indicator variable*

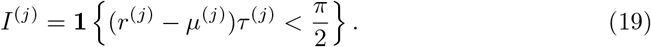

*Then*

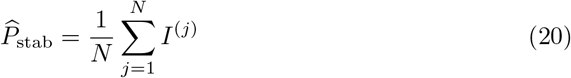

*is an unbiased estimator of P*_stab_.

*Proof*. Each *I*^(*j*)^ is a Bernoulli random variable with success probability

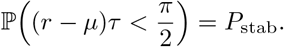

By linearity of expectation,

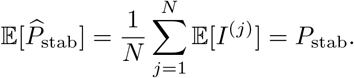

#### Proposition 3

**(Sampling distribution of the estimator)** *Under the assumptions above*,

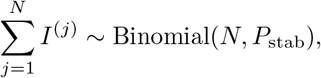

*and therefore*

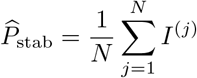

*Satisfies*

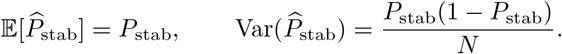

*Moreover, by the central limit theorem*,

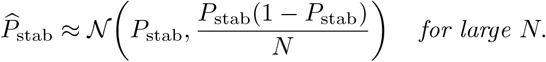

An asymptotic 95% confidence interval is therefore given by

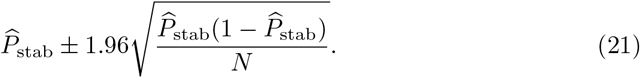

Independence is assumed across Monte Carlo samples, but not within the parameter vector (*r, µ, τ*). In particular, the joint distribution may encode ecological dependencies between recruitment, mortality, and delay.

To improve sampling efficiency, Latin Hypercube sampling (LHS) can be used in place of naive Monte Carlo. Let *F*_*r*_, *F*_*µ*_, and *F*_*τ*_ denote the marginal distribution functions. LHS partitions the unit interval into *N* strata for each parameter, samples once from each stratum, and combines the samples via random permutation before applying inverse transforms. The resulting design improves space-filling properties while preserving unbiased estimation of *P*_stab_.

### Numerical procedures

The delayed system (1) is integrated numerically using the method of steps, which converts the delay differential equation into a sequence of ordinary differential equations on successive intervals of length *τ* . Unless otherwise stated, a constant history function *x*(*t*) = *φ*(*t*) ≡*x*\* is used for *t* ∈ [−*τ*, 0].

To ensure reproducibility and clarity, the computational procedure used to estimate the probability of stability *P*_stab_ is summarized below.

#### Algorithm 1

Monte Carlo estimation of the probability of stability *P*_stab_

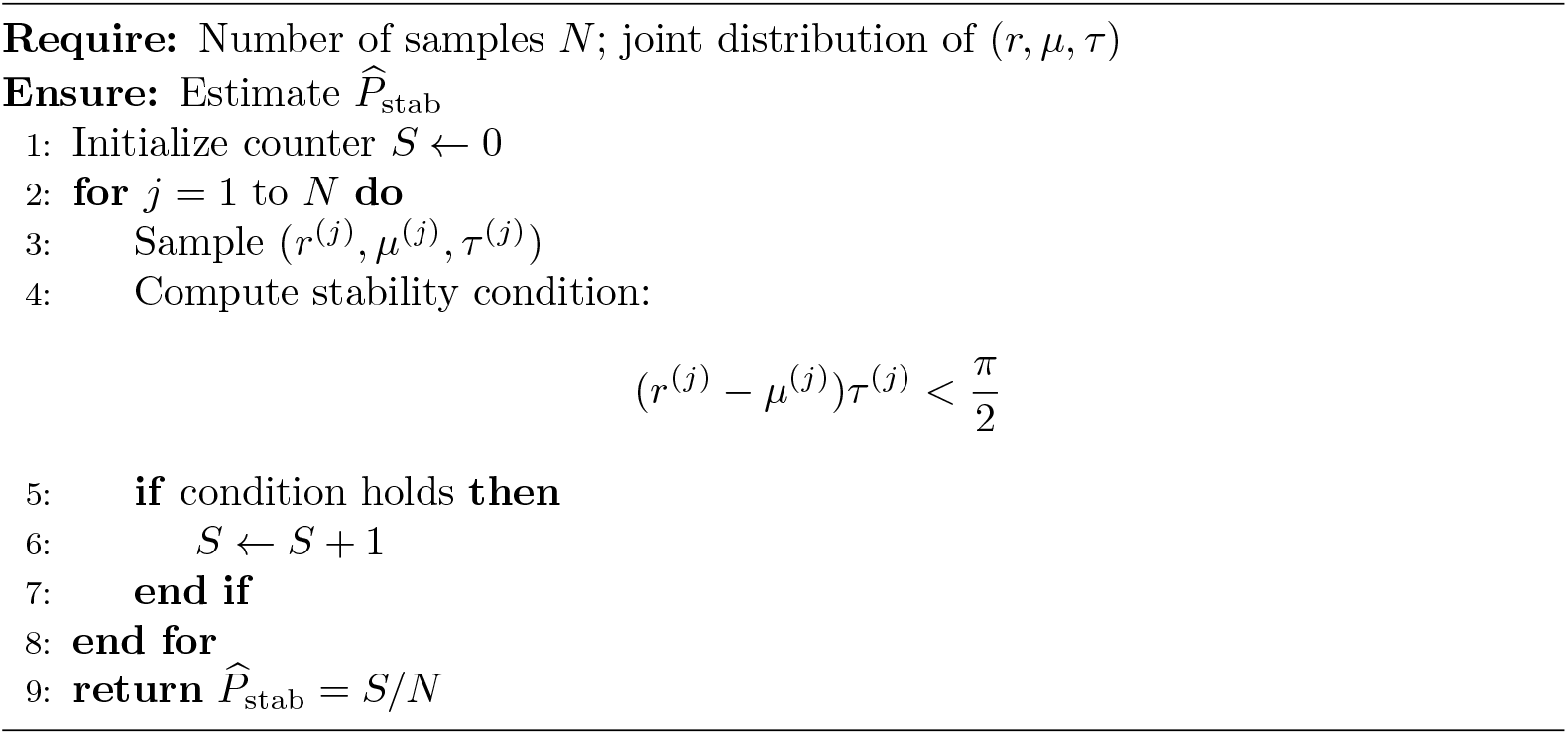

For improved sampling efficiency, Latin Hypercube sampling (LHS) can be used in place of standard Monte Carlo by replacing the sampling step in Algorithm 1 with a stratified sampling scheme based on the marginal distributions of *r, µ*, and *τ* .

To illustrate the theoretical results, we consider:

1. time-domain simulations in stable, near-critical, and unstable regimes;
2. scenario-based estimation of *P*_stab_ under parameter uncertainty;
3. heat-map visualization of *P*_stab_ across parameter combinations.

### Parameter ranges and biological interpretation

The parameter ranges employed in the probabilistic and sensitivity analyses are intended to represent exploratory ecological regimes rather than a single site-specific calibration. The objective is to examine how plausible variation in mosquito demographic processes modifies the stability of the resulting transmission potential, and to identify which ecological mechanisms are the primary drivers of stable or unstable regimes.

The recruitment parameter *r* is interpreted as an effective demographic rate summarizing the combined contribution of oviposition, immature survival, and maturation into the adult female mosquito population. In this reduced framework, *r* is not a directly observed field quantity, but rather a composite ecological growth parameter reflecting variability in larval habitat productivity and reproductive success across different landscapes. Variations in *r* capture differences in the quality and availability of aquatic breeding sites, which strongly influence the demographic pressure acting on the stability threshold.

The mortality parameter *µ* represents adult mosquito mortality and is the most directly interpretable parameter from a biological and operational perspective. It reflects variation in adult survival across ecological conditions and vector control settings, driven by environmental stress, predation, and intervention measures. Changes in *µ* are therefore indicative of the efficacy of adult-targeted vector control tools, such as long-lasting insecticide-treated nets and indoor residual spraying. In the context of stability, *µ* acts as a moderating force that counterbalances the destabilizing effects of rapid recruitment and delayed feedback.

The delay parameter *τ* represents the effective developmental and density-dependent feedback delay. Rather than being interpreted strictly as the egg-to-adult development time, *τ* captures the time required for past cohort structure and density-dependent processes (e.g. larval crowding) to influence current adult abundance. As such, *τ* defines the depth of biological memory in the system. Although mosquito development is known to vary with environmental conditions such as temperature and nutrition, we adopt a time-invariant approximation in this baseline analysis in order to isolate the qualitative effects of temporal memory on population regulation.

The numerical ranges used for these parameters, together with the alternative scenarios considered in the probabilistic analysis, are summarized in Table 2. These ranges span contrasting ecological regimes, from low to high recruitment, varying levels of adult mortality, and short to long developmental delays. They are not intended to reproduce a specific ecological setting, but rather to provide a transparent framework for exploring how demographic variability shapes stability.

**Table 1.** Exploratory parameter ranges used in the probabilistic and sensitivity analyses.

| Parameter | Baseline range | Alternative scenario ranges | References |
| --- | --- | --- | --- |
| $r$ | [0.12, 0.50] | Slower recruitment: [0.10, 0.22];<br>Faster recruitment: [0.30, 0.60] | [3, 33, 34] |
| $\mu$ | [0.05, 0.15] | Lower mortality: [0.02, 0.08];<br>Higher mortality: [0.10, 0.20] | [2, 3, 25, 35, 36] |
| $\tau$ | [5, 20] | Shorter delay: [2, 8]; Longer delay: [12, 25] | [34, 37, 38] |

**Table 2.** Estimated stability probability *P*_stab_ across illustrative parameter scenarios.

| Scenario | r range | μ range | τ range | MC P-hat_{stab} | LHS P-hat_{stab} |
| --- | --- | --- | --- | --- | --- |
| Baseline | [0.12, 0.50] | [0.05, 0.15] | [5, 20] | 0.3306 | 0.3318 |
| Shorter delays | [0.12, 0.50] | [0.05, 0.15] | [2, 8] | 0.7701 | 0.7798 |
| Longer delays | [0.12, 0.50] | [0.05, 0.15] | [12, 25] | 0.1818 | 0.1820 |
| Higher mortality | [0.12, 0.50] | [0.10, 0.20] | [5, 20] | 0.4614 | 0.4622 |
| Lower mortality | [0.12, 0.50] | [0.02, 0.08] | [5, 20] | 0.2018 | 0.2034 |
| Slower recruitment | [0.10, 0.22] | [0.05, 0.15] | [5, 20] | 0.8800 | 0.8812 |
| Faster recruitment | [0.30, 0.60] | [0.05, 0.15] | [5, 20] | 0.0335 | 0.0368 |

Within this framework, recruitment-related variation reflects differences in larval habitat productivity, mortality captures the impact of adult-targeted interventions, and delay represents environmentally driven changes in mosquito development and feedback processes. By quantifying the transition between stable and unstable regimes across these ranges, the partial rank correlation coefficient (PRCC) analysis provides a direct link between the mathematical stability condition and the identification of key ecological drivers. This perspective allows us to assess not only the level of transmission potential, but also its dynamical stability and sensitivity to ecological variability, thereby supporting the prioritization of vector control strategies.

## Results

### Dynamic regimes across the delay threshold

The system exhibits a clear transition in dynamical behavior at the threshold

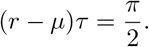

When (*r* − *µ*)*τ < π/*2, perturbations decay and the system converges toward equilibrium, either monotonically or through damped oscillations. As the threshold is approached, oscillations decay increasingly slowly, indicating the onset of a near-critical regime characterized by long transient dynamics and critical slowing down.

When (*r* −*µ*)*τ > π/*2, the equilibrium loses stability and the system generates sustained oscillations. These correspond to self-maintained limit-cycle-like dynamics driven purely by delayed demographic feedback.

Because *R*_0_(*t*) = *C*_0_*x*(*t*), the transmission potential inherits these qualitative behaviors directly (Fig. 2). As a result, delayed mosquito demography can generate autonomous oscillations in transmission potential, even in the absence of environmental forcing.

**Fig 2.**
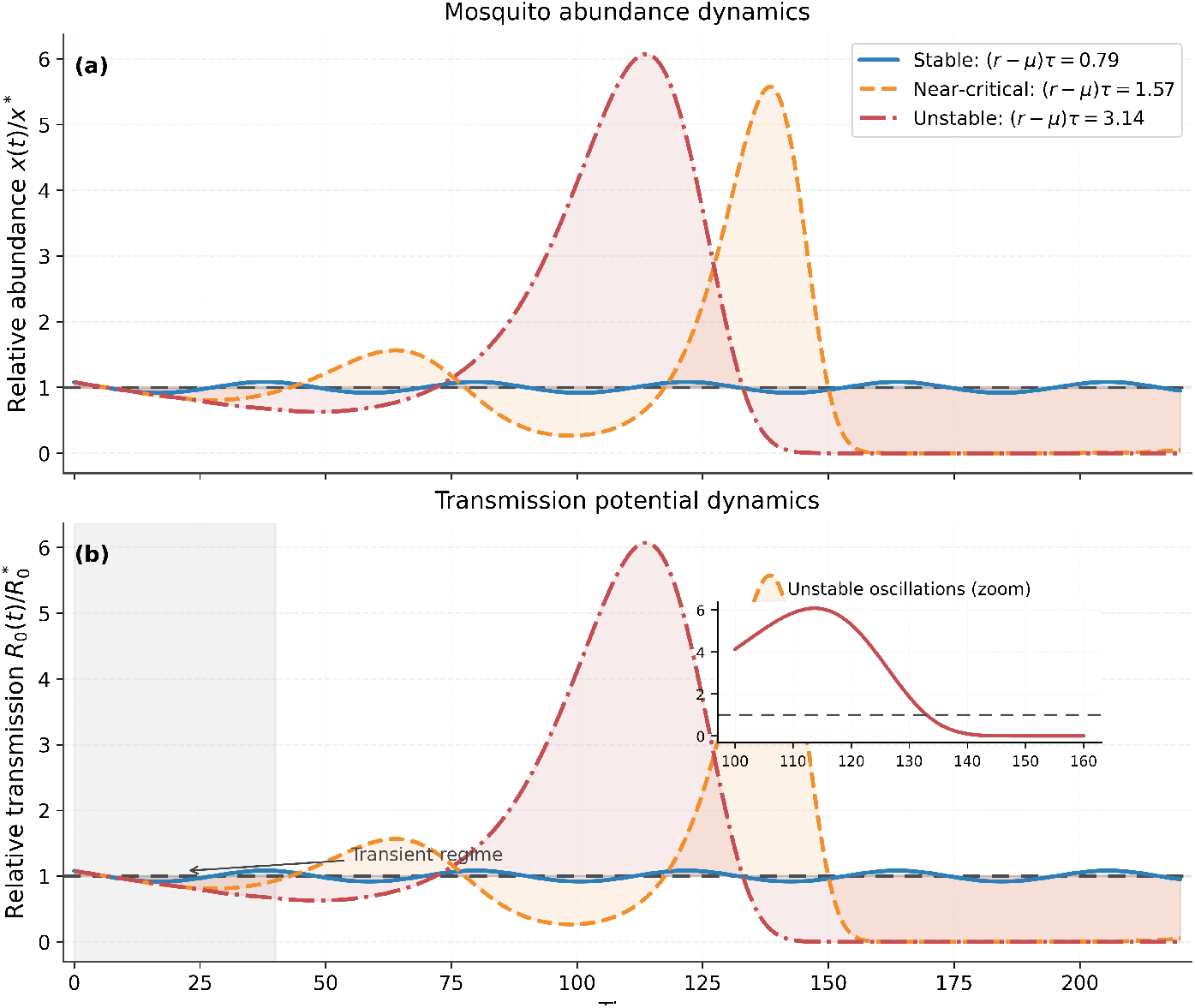
Delay-driven dynamical regimes in the mosquito population and the associated transmission potential. The top panel (a) shows the normalized mosquito abundance *x*(*t*)*/x*\*, while the bottom panel (b) shows the normalized transmission potential *R*_0_(*t*)*/R*_0_*, under stable, near-critical, and unstable delay regimes. The stable regime corresponds to (*r* −*µ*)*τ < π/*2, the near-critical regime to (*r* −*µ*)*τ* ≈ *π/*2, and the unstable regime to (*r* −*µ*)*τ > π/*2. Dashed horizontal lines indicate the equilibrium level. The inset in panel (b) highlights the oscillatory amplification in the unstable regime, while the shaded region indicates the transient adjustment phase following initial perturbation. Together, the two panels demonstrate that the transmission potential inherits the qualitative delay-driven dynamics of mosquito abundance.

This demonstrates that the delay parameter *τ*, representing biological memory, is not merely a temporal shift but a structural driver of system dynamics. Temporal variability in malaria risk may therefore arise endogenously, rather than being driven solely by seasonal or external factors.

In particular, near the threshold, the system exhibits slow recovery and persistent oscillations, defining a vulnerable regime in which intrinsic dynamics may be difficult to distinguish from externally driven transmission peaks.

### Scenario-based probabilistic stability

Across all scenarios, the estimated stability probability varied widely, revealing a strong dependence on ecological conditions (Table 2).

Under baseline parameter ranges, the stability probability was

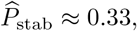

indicating that the system lies predominantly in an instability-prone regime, with delayed mosquito dynamics more likely to produce oscillatory or unstable behavior than stable convergence.

Parameter variation produced consistent and interpretable shifts. Reducing the delay substantially increased stability to approximately 0.77, whereas increasing the delay reduced it to approximately 0.18. This confirms the central role of developmental delay as the mechanism driving instability through biological memory.

Changes in mortality and recruitment further modulate this effect. Increasing adult mortality increased stability (≈ 0.46), while lower mortality reduced it (≈ 0.20).

Recruitment had the strongest modulating effect: slower recruitment produced a highly stable regime (≈0.88), whereas faster recruitment reduced stability to near zero (≈ 0.03). These results indicate that highly productive environments are particularly prone to instability when combined with delayed feedback.

Taken together, these results define a clear ecological gradient structured by the interaction between demographic rates and delay. Stability is favored by short delays, low recruitment, and high mortality, while instability emerges when delays are long and recruitment is rapid. The baseline scenario lies near this transition region, indicating that moderate environmental variability can shift the system between stable and unstable regimes.

From a control perspective, this gradient defines distinct stability regimes.

Environments characterized by prolonged developmental delays and high recruitment lie in an instability-prone zone, where transmission potential is intrinsically variable and difficult to predict. In contrast, increased adult mortality and reduced recruitment shift the system toward more stable and predictable dynamics. This highlights that vector control interventions may influence not only the magnitude of transmission, but also its dynamical stability and predictability under ecological variability.

From a control perspective, these patterns define a risk gradient across ecological regimes: productive habitats with rapid recruitment and long effective delays lie closer to unstable zones, whereas settings with higher adult mortality or slower recruitment remain in safer and more predictable transmission regimes.

### Probabilistic stability across parameter space

The heat maps reveal a clear and continuous transition between stable and unstable regimes across parameter combinations (Fig. 3).

**Fig 3.**
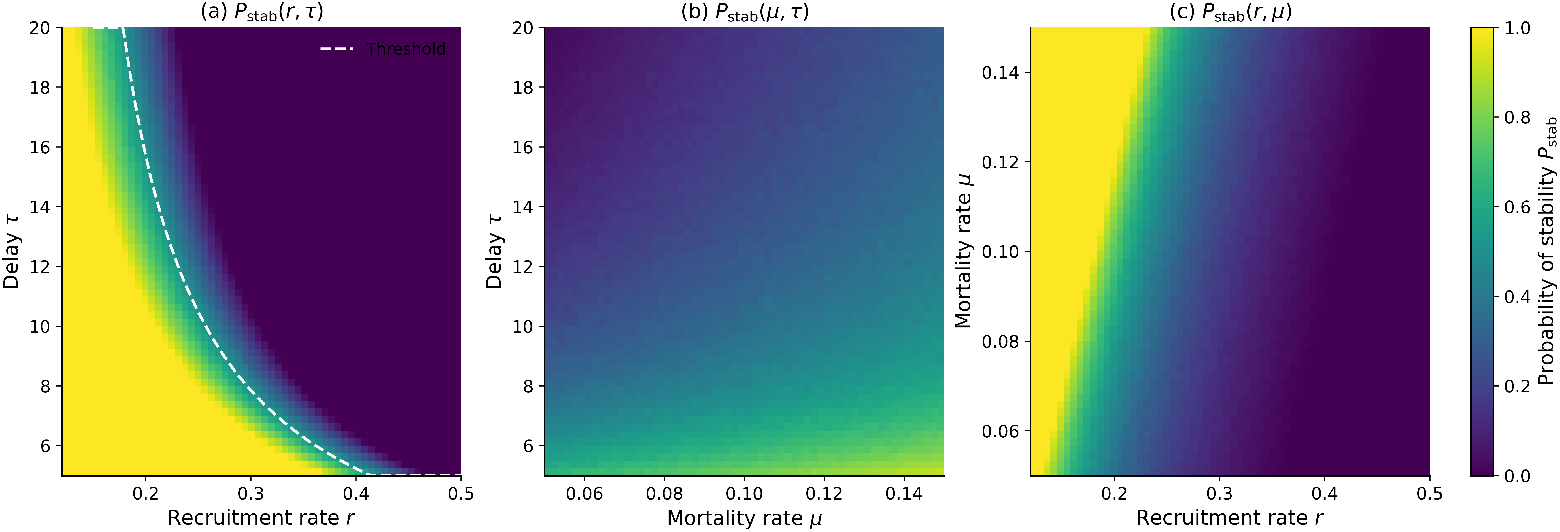
Three-panel heat-map representation of the probability of stability *P*_stab_. Panel (a) shows the dependence of stability on recruitment rate *r* and delay *τ*, panel (b) shows the dependence on mortality *µ* and delay *τ*, and panel (c) shows the dependence on recruitment and mortality. Color intensity represents the probability of stability. The dashed curve in panel (a) indicates the theoretical threshold (*r* − *µ*)*τ* = *π/*2, highlighting the transition between stable and unstable regimes.

Panel (a) shows that stability decreases sharply as both recruitment *r* and delay *τ* increase. In particular, even moderate increases in delay rapidly push the system toward instability when recruitment is high, highlighting the central role of developmental delay as the mechanism through which instability emerges.

Panel (b) shows that increasing mortality *µ* expands the stable region, partially counteracting the destabilizing effect of delay. However, this stabilizing influence is limited when delays are long, indicating that biological memory can dominate mortality effects in shaping system behavior.

Panel (c) confirms that recruitment strongly modulates stability, with high recruitment driving instability even under moderate mortality.

Across all panels, a transition band separates stable and unstable regimes. This band aligns closely with the theoretical threshold

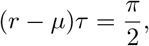

but appears broadened due to parameter uncertainty, forming a probabilistic rather than deterministic boundary.

These results define a “vulnerable zone” in parameter space where small changes in ecological conditions can shift the system between stable and unstable dynamics. This zone corresponds to conditions under which delayed feedback is sufficiently strong to destabilize the system, but not yet dominant enough to produce fully developed oscillations.

More broadly, the heat maps demonstrate that delay transforms a sharp theoretical threshold into a probabilistic regime structure. This provides a quantitative framework for assessing not only whether transmission is stable, but also how sensitive it is to ecological variability and environmental change.

This instability-prone region represents a critical zone for intervention planning, where small environmental or control-induced changes may lead to disproportionately large shifts in transmission dynamics.

### Global sensitivity of delay-induced stability

Global sensitivity analysis reveals a clear hierarchy in the drivers of stability, while highlighting the distinct roles of recruitment and delay in shaping transmission dynamics.

Recruitment exhibits the strongest destabilizing effect (PRCC_*r*_ ≈ −0.96), followed closely by the developmental delay (PRCC_*τ*_ ≈ −0.90). In contrast, mortality has a stabilizing effect (PRCC_*µ*_ ≈ 0.64). These results are consistent across sample sizes (Table 3), indicating a robust sensitivity ranking.

**Table 3.**
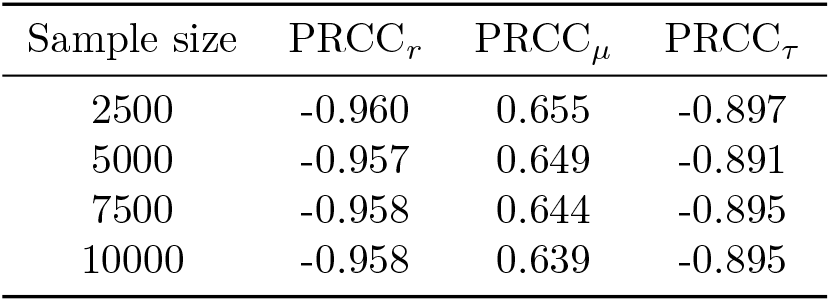
Convergence of PRCC values for the stability margin across increasing sample sizes.

| Sample size | $\text{PRCC}_r$ | $\text{PRCC}_\mu$ | $\text{PRCC}_\tau$ |
| --- | --- | --- | --- |
| 2500 | -0.960 | 0.655 | -0.897 |
| 5000 | -0.957 | 0.649 | -0.891 |
| 7500 | -0.958 | 0.644 | -0.895 |
| 10000 | -0.958 | 0.639 | -0.895 |

Importantly, the roles of recruitment and delay are complementary rather than interchangeable. The delay parameter *τ* represents the biological memory of the system and is the key mechanism responsible for the emergence of oscillatory and unstable dynamics. In the absence of delay, such instabilities cannot occur. By contrast, the recruitment rate *r* controls the intensity of this delayed feedback, amplifying or dampening its effect.

This distinction is reflected in the analytical stability condition (*r* − *µ*)*τ < π/*2, where instability arises from the interaction between the strength of population growth and the length of the memory timescale. The high sensitivity of the system to both *r* and *τ* therefore, confirms that delay-driven feedback is central to the dynamics, with recruitment acting as a dominant modulator.

From a biological perspective, this implies that environmental factors affecting developmental time (e.g. temperature, larval crowding) play a critical role in determining whether transmission dynamics become oscillatory, while factors affecting recruitment determine how strongly such oscillations are expressed. Together, these processes define the stability structure of malaria transmission under ecological variability.

## Discussion

The fundamental contribution of this study is the formalization of a shift in transmission theory: from the interpretation of *R*_0_ as a static threshold to its characterization as a dynamic, time-dependent object *R*_0_(*t*). By incorporating biological memory into the Ross–Macdonald framework, we demonstrate that the stability of malaria transmission is not merely a consequence of vector suppression, but an emergent property of demographic history. Our analysis reveals that stability is governed by the dimensionless quantity (*r* − *µ*)*τ* = *π/*2. When delayed feedback becomes sufficiently strong, the system transitions from convergence to intrinsic oscillatory or unstable regimes, even in the absence of explicit seasonal forcing. This suggests that the temporal variability frequently observed in many field data [23, 27, 39–41], often attributed solely to external environmental factors, may in part arise from intrinsic instability generated by delayed vector population processes. In this sense, *R*_0_(*t*) serves as a dynamical signature of the vector population’s demographic health.

A second central result is the recasting of the deterministic threshold into the probabilistic stability metric: *P*_stab_ = *P* ((*r*−*µ*)*τ < π/*2). In practice, ecological rates are rarely observed without error and vary significantly across sites, seasons, and intervention contexts [42, 43]. Under such uncertainty, a single threshold value is less informative than a stability index that quantifies the likelihood of remaining in a robust transmission regime [26, 44]. We propose that *P*_stab_ be interpreted not only as a mathematical probability, but as a decision-oriented transmission risk indicator. Low values indicate greater potential for unstable or difficult-to-predict transmission dynamics, whereas high values indicate more stable regimes. This index allows for the identification of settings in which transmission is not only high, but dynamically fragile, supporting the use of stability-based metrics as complementary tools for intervention prioritization and adaptive malaria control.

The PRCC analysis clarifies the biological drivers of this stability, showing that recruitment *r* and developmental delay *τ* are the dominant drivers of instability (PRCC_*r*_ ≈− 0.96, PRCC_*τ*_ ≈ −0.90). Importantly, these roles are complementary rather than interchangeable. The delay parameter *τ* represents the biological memory of the system and is the key mechanism that enables the emergence of oscillatory and unstable dynamics; in the absence of delay, such instabilities cannot occur. By contrast, the recruitment rate *r* controls the intensity of this delayed feedback, amplifying or dampening its effect.

This provides a clear link between ecological processes and transmission structure: recruitment and developmental delay are strongly shaped by larval habitat productivity and temperature, whereas mortality is more directly affected by adult-targeted interventions. This ranking suggests that interventions targeting larval habitat productivity and developmental dynamics, such as Larval Source Management (LSM), oviposition trapping, or strategies that reduce emergence, may have a stronger impact on stabilizing transmission potential than strategies acting solely on adult mortality. This aligns with broader calls to diversify the malaria vector-control toolbox and target a wider range of mosquito life-history processes [45].This is consistent with the analytical stability condition (*r* − *µ*)*τ* = *π/*2, which shows that instability arises from the interaction between the strength of population growth and the memory timescale, confirming the central role of delay in shaping transmission dynamics.

As global temperatures and rainfall patterns shift, vector parameters respond in tandem. Warmer conditions typically shorten maturation delays *τ* while accelerating recruitment *r* [42, 46–48]. *While these effects appear potentially offsetting in the (r* − *µ*)*τ* threshold, the acceleration of recruitment in high-productivity zones often pushes the system toward the unstable regime (*P*_stab_*→* 0). Consequently, climate change may increase the dynamical fragility of transmission. As stable regimes shift toward “near-critical” zones, the system becomes hypersensitive to transient environmental “shocks,” such as heatwaves or sudden habitat clearing [49–51]. Under these conditions, the transmission potential may exhibit prolonged “ringing” (slowly decaying oscillations), making the timing of interventions, and the interpretation of their success, increasingly difficult. An observed spike in transmission may be an endogenous echo of a past ecological shock rather than a failure of current control strategies.

A critical operational issue highlighted by our model is intervention lag. Because the vector population inherits its current state from a developmental window [*t* − *τ, t*] the effect of an intervention on *R*_0_(*t*) is not instantaneous. Changes in mosquito demography may require several maturation cycles before a full response is visible. This biological inertia implies that short-term post-intervention evaluations may systematically underestimate the long-term effect of control measures. This reinforces the importance of timing, not just for deployment, but for assessment, ensuring that efficacy is measured over a window that accounts for the propagation of feedback across successive cohorts.

Finally, our framework connects naturally to the future of disease intelligence. In heterogeneous landscapes, *P*_stab_ provides a more nuanced interpretation of intervention performance than mean transmission indicators alone. This is closely aligned with current efforts to use modeling for risk stratification and progressive disease control under uncertainty [52–58]. While developed in a malaria setting, this logic is applicable to other vector-borne diseases where ecological uncertainty and delayed vector processes shape outbreak risk.

The present model is intentionally minimal, acting as a baseline framework to isolate the role of mosquito demographic memory. It does not yet include seasonality, distributed delays, or a fully coupled host-vector system. This simplicity allows for a transparent analysis of the mechanism before moving to more elaborate formulations. Future extensions will prioritize: 1) Coupling the framework to seasonal drivers where temperature and rainfall alter *r, µ* and *τ* simultaneously, 2) Combining *P*_stab_ with explicit intervention models to evaluate how specific tools move a system across regime diagrams and 3) Embedding the framework in spatially explicit models to map stability probability across landscapes alongside other risk indicators.

## Conclusion

This study provides a transparent, delay-based extension of the Ross–Macdonald framework that transforms our understanding of malaria transmission from a static threshold into a dynamic, uncertainty-aware system. We established that the stability of the transmission potential *R*_0_(*t*) is governed by the dimensionless quantity (*r* − *µ*)*τ*, identifying a critical threshold at 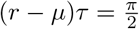 . By recasting this boundary as the probability of stability, 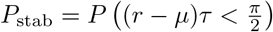, we offer a new index for risk stratification that accounts for ecological variability. Ultimately, our work suggests that the most effective vector control strategies are those that not only suppress the intensity of transmission but also shift the system toward more predictable and stable regimes, providing a robust foundation for decision-making in a changing environment.

## Code availability

Python scripts used for numerical simulations, probabilistic analyses, and supplementary verification are provided in the Supporting Information and will be made publicly available through a GitHub repository upon publication.

## Data availability

This study does not use empirical datasets. All numerical simulations and analyses are reproducible using the Python scripts provided in the Supporting Information.

## Supporting information

**S1 Fig. Delay-driven dynamical regimes in mosquito abundance and transmission potential**. Supplementary time-domain simulations illustrating stable, near-critical, and unstable regimes generated by delayed mosquito demographic feedback. The figure presents the corresponding dynamics of mosquito abundance *x*(*t*) and transmission potential *R*_0_(*t*) across the stability threshold (*r* −*µ*)*τ* = *π/*2. Simulations were performed using the method of steps with an explicit Euler discretization and a fixed delay. The apparent smoothness of the trajectories results from the use of a small numerical time step and high-resolution rendering rather than post-simulation smoothing.

**S2 Fig. Characteristic-root crossing near the delay-induced stability threshold**. Supplementary visualization of the dominant characteristic roots of the delay differential equation obtained using the Lambert *W* representation of the characteristic equation. The figure illustrates how the dominant roots move across the imaginary axis as the dimensionless quantity (*r* − *µ*)*τ* increases, confirming the analytically derived threshold (*r* − *µ*)*τ* = *π/*2. Additional complex-plane snapshots illustrate the distribution of characteristic roots on the stable side, at the critical threshold, and in the unstable regime.

**S3 Fig. Partial Rank Correlation Coefficient (PRCC) sensitivity analysis**. Supplementary global sensitivity analysis showing the PRCC values and bootstrap confidence intervals associated with the stability margin

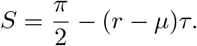

The analysis quantifies the relative contribution of recruitment, mortality, and developmental delay to delay-induced instability in the transmission potential.

**S1 Table. PRCC convergence across sample sizes**. Convergence of the Partial Rank Correlation Coefficients computed using Latin Hypercube sampling for increasing sample sizes (*N* = 2500, 5000, 7500, and 10000). The stability of the PRCC estimates across sample sizes demonstrates the robustness of the sensitivity ranking.

**S1 Appendix. Supplementary mathematical and numerical methods**. Detailed derivation of the characteristic-root formulation for the delayed mosquito demographic equation, including the Lambert *W* -based representation of characteristic roots and supplementary discussion of root-crossing behavior near the stability threshold. The appendix also provides additional details on the numerical implementation of the delay simulations, Monte Carlo estimation of *P*_stab_, heat-map construction, and the PRCC methodology.

**S1 File. Reproducible Python code and numerical analysis scripts**. Compressed archive containing all Python scripts used to generate the main and supplementary numerical results, including delay-regime simulations, probabilistic stability heat maps, PRCC sensitivity analysis, and characteristic-root verification analyses.

## Acknowledgments

AMGB, PB acknowledge support from the European Union’s Horizon Europe research and innovation programme under grant agreement n°101000467, acronym ‘‘IMPACTING” (Integrated Multi-vector-borne diseases Platform to Assess how global Change impacts Transmission using Innovative systems modeling, Novel monitoring tools, and transmission blockinG micro-organisms) “PB has acknowledge support from Open Philanthropy (SYMBIOVECTOR), the Bill and Melinda Gates Foundation (INV0225840). International Centre of Insect Physiology and Ecology (ICIPE) also receives funding and support from The Swedish International Development Cooperation Agency (Sida); the Swiss Agency for Development and Cooperation (SDC); the Australian Centre for International Agricultural Research (ACIAR); the Norwegian Agency for Development Cooperation (Norad); the German Federal Ministry for Economic Cooperation and Development (BMZ); and the Government of the Republic of Kenya. The views expressed herein do not necessarily reflect the official opinion of the donors.

